# Semi-automated protocol to quantify and characterize fluorescent three-dimensional vascular images

**DOI:** 10.1101/2022.05.05.490827

**Authors:** Danny F. Xie, Christian Crouzet, Krystal LoPresti, Yuke Wang, Christopher Robinson, William Jones, Fjolla Muqolli, Chuo Fang, David H. Cribbs, Mark Fisher, Bernard Choi

## Abstract

The microvasculature facilitates gas exchange, provides nutrients to cells, and regulates blood flow in response to stimuli. Vascular abnormalities are an indicator of pathology for various conditions, such as compromised vessel integrity in small vessel disease and angiogenesis in tumors. Traditional immunohistochemistry enables visualization of tissue cross-sections containing exogenously labeled vasculature. Although this approach can be utilized to quantify vascular changes within small fields-of-view, it is not a practical way to study the vasculature on the scale of whole organs. Three-dimensional (3D) imaging presents a more appropriate method to visualize the vascular architecture in tissue. Here we describe the complete protocol that we use to characterize the vasculature of different organs in mice encompassing the methods to fluorescently label vessels, optically clear tissue, collect 3D vascular images, and quantify these vascular images with a semi-automated approach. To validate the automated segmentation of vascular images, one user manually segmented fifty random regions of interest across different vascular images. The automated segmentation results had an average sensitivity of 80±8% and an average specificity of 90±5% when compared to manual segmentation. Applying this procedure of image analysis presents a method to reliably quantify and characterize vascular networks in a timely fashion. This procedure is also applicable to other methods of tissue clearing and vascular labels that generate 3D images of microvasculature.

## Introduction

The microvasculature facilitates gas exchange, provides nutrients to cells, and regulates blood flow in response to stimuli. Thus, it plays a fundamental role in the survival and health of tissues and organs. Vascular abnormalities are an indicator of pathology for various conditions, such as compromised vessel integrity in small vessel disease and angiogenesis in tumors (1,2). In addition, impaired blood flow to the brain is associated with neurodegenerative diseases such as Alzheimer’s disease (3).

Traditional immunohistochemistry enables visualization of tissue cross-sections containing exogenously labeled vasculature. Although this approach can be used to quantify vascular changes within small fields-of-view, it is not a practical way to study the vasculature on the scale of whole organs. Furthermore, traditional immunohistochemistry requires the sectioning of tissue into thin (6-40 μm) sections, which effectively limits visualization of features to planar views and thus impedes facile 3D visualization of vascular architecture.

Volumetric imaging of tissue samples is vital to studying the microvasculature in its native state. The main limitation to 3D imaging of tissue samples is optical scattering resulting from the microscopic variations of refractive index occurring in most biological tissues. Organic solvents and aqueous solvents have been used to reduce tissue turbidity by achieving refractive index matching (4,5). CLARITY is a popular method used for tissue clearing that involves embedding of the sample into a hydrogel and using an electric current to remove lipids from the sample (6). Other tissue clearing procedures have advantages and disadvantages associated with clearing time, extent of clearing, and preservation of original tissue characteristics (size, endogenous fluorescence, etc.).

Quantitative analysis of microvasculature is essential to understand how structural variations in the microvascular network may change for different pathological states (7). The feasibility of such analysis depends on accurate, ideally automated segmentation methods for isolating the microvasculature from the background due to the hundreds of gigabytes of data generated with high-resolution 3D imaging. Various groups have developed automated algorithms to perform quantitative characterization of vascular images (8,9). Each method varies in complexity, processing time, and computational requirements.

We currently use iDISCO (immunolabeling-enabled three-dimensional imaging of solvent-cleared organs) as our primary method of tissue clearing (4,10). Briefly, iDISCO consists of methanol dehydration, lipid removal with dichloromethane, and refractive index matching with dibenzyl ether. iDISCO is a fast, simple-to-implement clearing method that enables deep-tissue imaging. It is compatible with many exogenous labels often used for traditional immunohistochemistry. We previously demonstrated the effectiveness of iDISCO in combination with lectin-DyLight-649 for 3D visualization of vasculature in a mouse brain (10). We also have used Prussian blue labelling of hemosiderin (11–13), a by-product of cerebral microhemorrhages, with iDISCO-cleared brains and lectin-DyLight-649 labelling.

In this paper, we first review different methods for vascular labeling in conjunction with optical clearing that have been published by other groups. Next, we describe the complete protocol that we use for imaging the vasculature in different organs in mice. Specifically, we report on sample collection and perfusion of lectin-DyLight-649 followed by application of additional labels as desired and optical clearing. We then describe our procedure to obtain 3D confocal microscopy images. Finally, we describe our semi-automated approach to process the resulting images and quantify the vascular architecture in three dimensions.

### Vascular imaging with various vessel labels and optical clearing techniques

Through reviewing the literature on 3D vascular imaging of cleared tissue, one will quickly realize the large variability in methodology across different experiments (see Table 1 for specific references). For example, blood vessels can be visualized with transgenic animals expressing a fluorescent protein in endothelial cells, labeled *in vivo* with intravascular dyes, or labeled *ex vivo* with immersion-based antibody labelling. In addition, tissue preparation can vary from thick tissue sections and whole organs to whole-mount specimens, depending on the purpose of an experiment. Finally, clearing procedures can be grouped into aqueous solvents (14–19), organic solvents, and hydrogel-based.

**Table 1.** Summary of different approaches to visualize 3D microvasculature in optically cleared tissues.

| Clearing agent(s) | Organ(s) | Vascular label | Quantification Method/Tool | Quantified Metrics |
| --- | --- | --- | --- | --- |
| MACS (14) | brain, heart, lung, spleen, femur, kidney, spinal cord, embryo, entire body | Dil, tetramethylrhodamine-conjugated dextran | Custom algorithm via Python, Imaris: surface function | glomerulus number and glomerulus volume; general metrics (imaging depth, size change, fluorescence intensity) |
| SeeDB, SeeDB2, 3DISCO, uDISCO, iDISCO, CUBIC, simplified CLARITY method, 75% v/v glycerol, Ce3D, FRUIT (23) | lymph nodes | CD31 | Imaris: surface function, ImageJ | general metrics (imaging depth, size change, fluorescence intensity) |
| sodium dodecyl sulfate/sodium deoxycholate with ScaleCUBIC-2 (24) | brain | RITC-Dex-GMA, Texas red lectin, CD31, αSMA | N/A | capillary diameters, signal-to-noise ratio |
| iDISCO (4) | brain, peripheral nervous system, kidney, muscle, heart, whisker pad, entire embryo | PeCAM | N/A | N/A |
| iDISCO with CUBIC (25) | ovary | endomucin | N/A | N/A |
| FDISCO (20) | brain, kidney | AlexaFluor647, CD31, lectin-DyLight-649 | Imaris, ImageJ | general metrics (size change, imaging depth, fluorescence quantification) |
| ScaleS (15) | brain | Texas Red lectin | custom C++, commercial Volocity, commercial Igor Pro | microglia and Aβ plaques with custom C++, distance between microglia and Aβ plaques |
| 3DISCO (26) | brain | FITC albumin-gelatin hydrogel | Imaris, ImageJ | vessel density, vessel diameter |
| thiazone with PEG-400 (27) | dorsal skin | N/A (imaged blood flow via LSI) | N/A | bloodflow via LSCI |
| iDISCO (28) | brain | Dye-conjugated secondary antibodies | ClearMap | Aβ deposits |
| CLARITY, TDE (29) | brain | lectin-FITC, gel-BSA-FITC, gel-BSA-TRITC | Markov random field based algorithm, ImageJ, Amira 5.3 software | automatic segmentation, vessel diameter, vessel length |
| FACT (30) | brain, spinal cord, heart, lung, adrenal gland, pancreas, liver, esophagus, duodenum, jejunum, ileum, muscle, bladder, ovary, uterus | CD31, auto fluorescence | Imaris | segmentation |
| CUBIC, BABB (16) | heart | FITC-lectin, 649-lectin, CD31 | N/A | N/A |
| BABB (31) | heart | PECAM1 | N/A | N/A |
| DBE, SCALE, CLARITY, CUBIC (32) | heart and embryo | GFP |  |  |
| vDISCO (33) | whole mouse | GFP, lectin | ImageJ, ClearMap | signal level, microglia distribution |
| 3DISCO (34) | brain | tomato lectin | N/A | N/A |
| CLARITY (35) | brain | fluorescein-conjugated tomato lectin | Amira | area, volume, perimeter, and length of stained vessel |
| CLARITY with ScaleA2 (36) | ovary, uterus, lung, liver | tdTomato | Ilastik, Imaris | segmentation, total vessel length, vessel mean diameter, vessel straightness, total number of branching points |
| PEGASOS (22) | whole mouse | αSMA, GS-IB4 isolectin dye, collagen IV | Imaris | vessel density |
| iDISCO (9) | brain | CD31, podocalyxin, collagen IV, smooth muscle actin, transgelin, von Willebrand factor | ClearMap 2.0, TubeMap | vascular density, vascular organization, vascular remodeling |
| ethyl-cinnamate (37) | liver, skin, lung, heart, muscles, pancreas, brain, kidney | In-house developed NIR fluorescent dye (MH148-PEI) | Leica LAS X | segmentation, volume of glomerulus |
| X-CLARITY (38) | placenta | Dil | Imaris, Image-Pro Premier | N/A |
| CLARITY, iDISCO (39) | brain | lectin-DyLight, lectin-FITC, anti-CD31 | Imaris, segmentation | number of branches, total vessel length, total vessel volume, total vessel area, diameter of single segments per volume, distance between cells and nearest vessel |
| PEGASOS (40) | bone, teeth | tdTomato | Imaris | blood vessel volume |
| ScaleS (17) | pancreas | lectin-DyLight | N/A | N/A |
| CLARITY (41) | retina | Griffonia lectin | Angiotool, ImageJ, Vaa3D, MATLAB, APP-2.0 | network tracing, vessel percentage area, total number of junctions, junction density, total vessel length, average vessel length, total number of endpoints, mean lacunarity |
| No clearing approach (42) | retina, heart | Dil | N/A | N/A |
| FocusClear (18) | small intestine | Dil | N/A | N/A |
| FocusClear, ScaleSQ, RIMS, sRIMS (19) | brain | Lectin-Dylight649 | MATLAB | optical properties, vascular density |
| FRUIT 100 (43) | brain tumor | Dil | N/A | N/A |
| FocusClear (44) | brain | Dil | N/A | optical properties |
| FocusClear (45) | brain | Dil | MATLAB | functional vascular density |

When selecting a clearing procedure, it is essential to consider its compatibility with the chosen vascular label. Endogenous fluorescence intensity is known to be reduced when using the iDISCO protocol; hence, an alternative vascular label would be preferred. Imaging of cleared samples is typically performed with confocal microscopy or light-sheet microscopy. Lastly, vascular-related metrics, such as vessel diameters and vessel density, are quantified with existing software such as FIJI and Imaris, or custom-designed software such as ClearMap 2.0.

We have reviewed several optical clearing methods utilized to image vasculature. They are organized in Table 1 by clearing method, tissue type, vessel label, and quantification. Vascular images in cleared tissue from select publications are included in Fig 1B-D. FDISCO is an organic solvent-based clearing method derived from the 3DISCO method. In developing FDISCO, Qi *et al*. found that they could achieve superior fluorescent signals of CD31-labelled vessels by optimizing temperature and pH conditions (Fig 1B, 1B’) (20,21). Moy *et al*. visualized vessels with an intravascular fluorescent dye along with FocusClear (Figure 1C, 1C’) (8). They were able to quantify functional vascular density in different regions of cardiac tissue with a custom MATLAB algorithm (21). With a polyethylene glycol-associated solvent system (PEGASOS), researchers developed a clearing technique that preserved both hard and soft tissue structures (Fig 1D, 1D’) (22), allowing for vascular imaging throughout an entire specimen.

**Fig 1.**
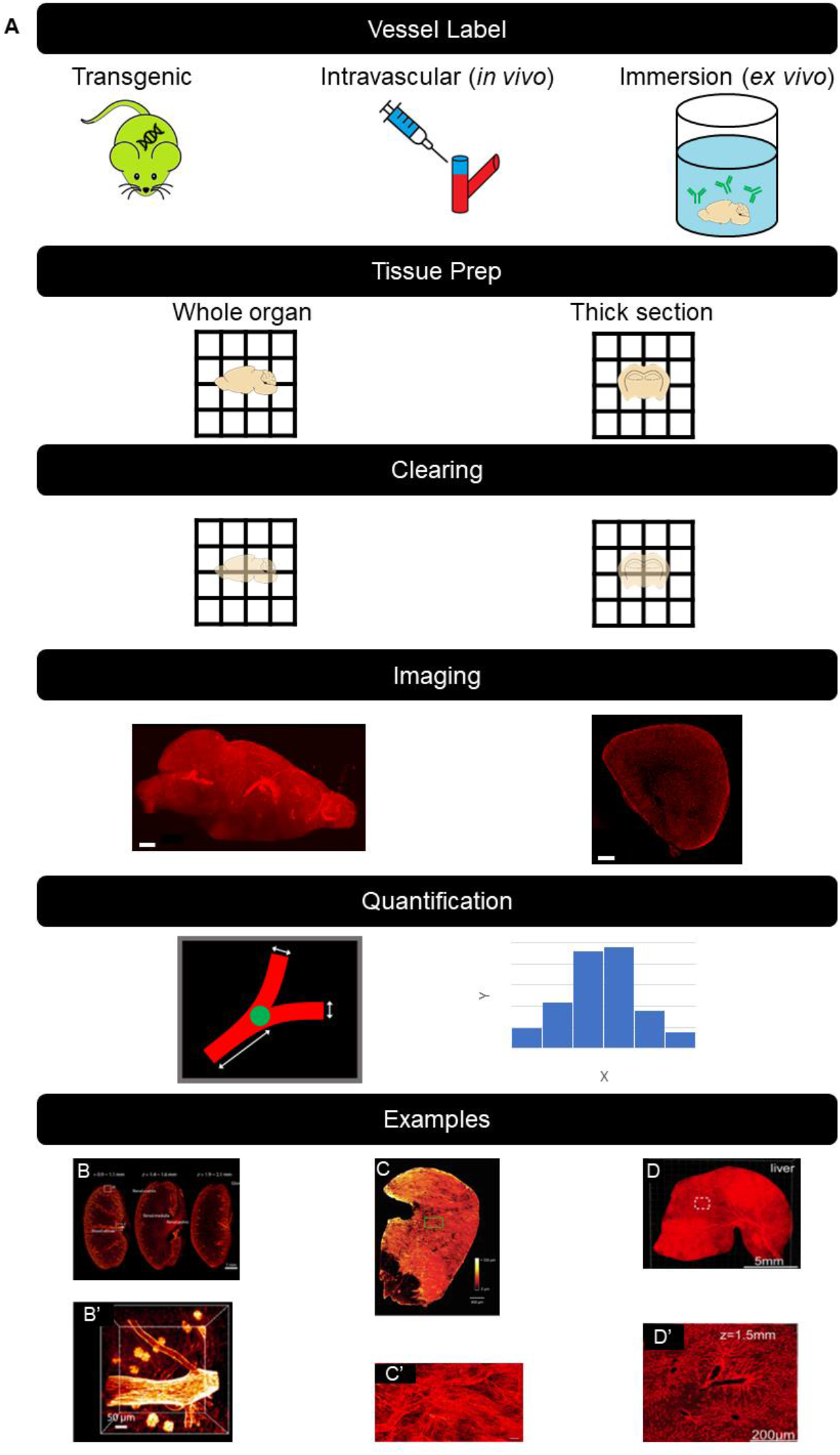
A) Illustrations of the critical steps and workflow for vascular visualization in cleared tissue. B-D) Examples of vascular visualization in literature. B) Visualization of CD31-labeled mouse kidney vasculature with FDISCO (20). B’) Magnified view of 1B. C) Visualization of Dil-labeled mouse heart vasculature with FocusClear (8). C’) Magnified view of 1C. D) Visualization of liver vasculature from *Tie2-Cre* mouse model with PEGASOS (22). D’) Magnified view of 1D).

### Three-dimensional quantitative analysis of microvasculature imaged in thick tissue sections

Microvascular labeling is performed with an injection of a lectin conjugated to a fluorophore. In our work, we have focused on the DyLight-649 fluorophore. As the lectin travels through the circulatory system, it binds to glycoproteins adjacent to endothelial cells of the vascular wall. This binding allows for labeling of the vascular network within every organ of the body.

An overview of our workflow is outlined below in Fig 2. We administer lectin-DyLight-649 via retroorbital injection. The lectin is allowed to circulate throughout the body prior to cardiac perfusion with saline followed by formalin. Mouse brains are then extracted, bisected into hemispheres, and sectioned into thick (0.5-1.0mm) sections. Each section is then imaged using confocal microscopy to generate image stacks of the complete section throughout its entire thickness. Segmentation is performed by a custom MATLAB (MathWorks, Natick, MA) script. Finally, neuTube, an open-source neuron tracing software, is used to extract diameter measurements of the vessel structures (46). Below, we describe in detail the materials and methods we use.

**Fig 2.**
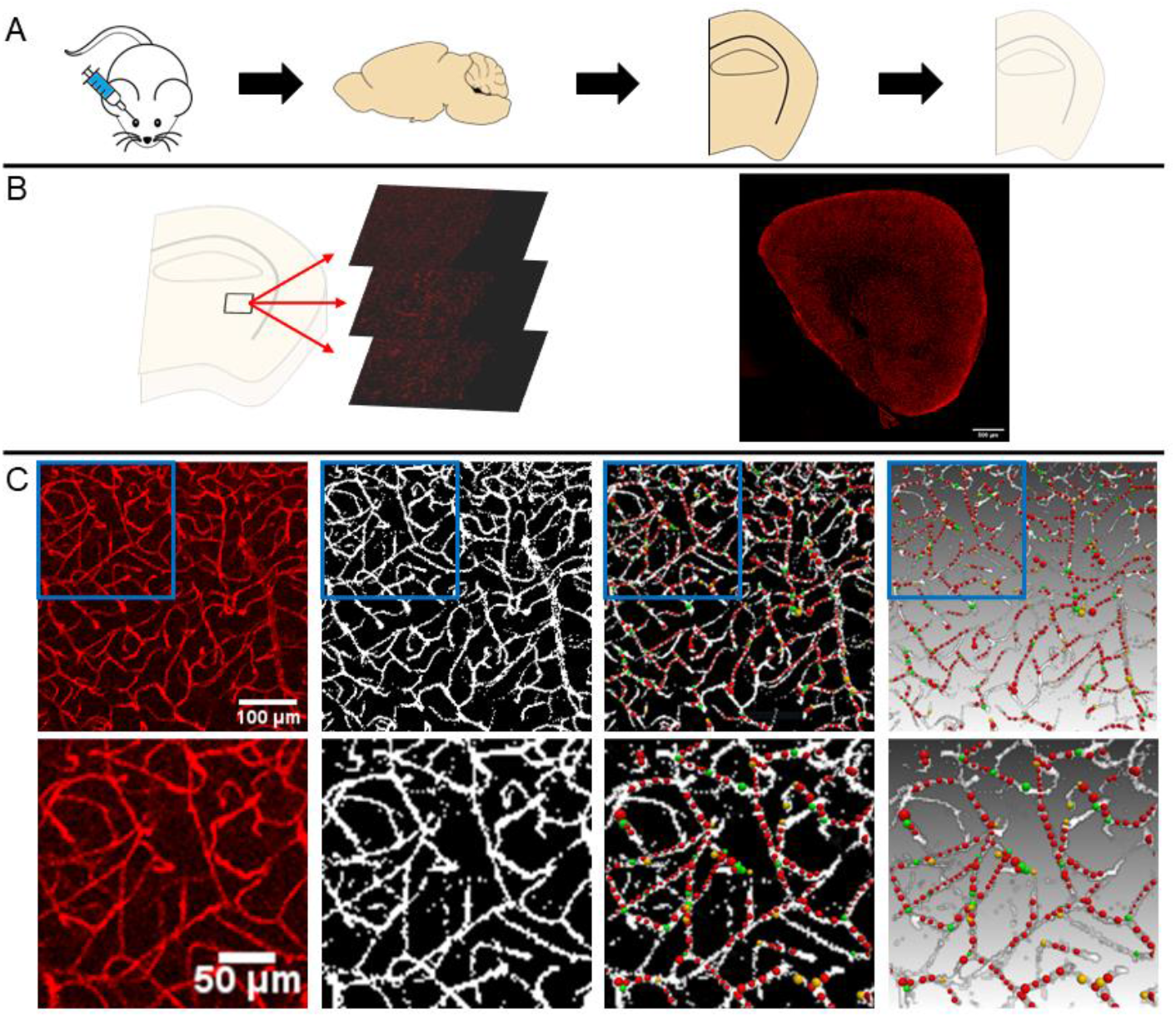
Workflow for 3D visualization and quantitation of microvasculature in thick tissue sections. A) Vasculature is labeled via retroorbital injection of lectin-DyLight-649. Brains are then sectioned and cleared. B) Depiction of z-stack imaging and tile stitching to visualize an entire sample in 3D via confocal microscopy. C) From left to right: representative example of a raw image of the vasculature, segmented vasculature via MATLAB, and traced vasculature via neuTube.

## Methods

### Equipment and supplies

#### Reagents

- Saline solution (Aspen Veterinary Resources, NDC No. 46066-807-50)
- 10% formalin solution (Sigma-Aldrich Product Code HT50-1-1)
- PBS with 0.02% sodium azide solution (Syringa Lab Supplies, Part No. 11001)
- Potassium ferrocyanide (Sigma: P3289-500G)
- Hydrochloric acid (Fisher: A144-212)
- Methanol (Fisher: A412S)
- Deionized water
- Dichloromethane (Sigma: 270997-100ML)
- Dibenzyl ether (Sigma: 108014-1KG)

#### Equipment

- 2x 10 mL syringes (Sigma-Aldrich: Z683604)
- 1x 23G x ¾ x 12” butterfly needle (Vaculet, 26766)
- 3x hemostat clamps (Excelta: 37SE)
- 1x bone scissors (Fine Science Tools: 91604-09)
- 1x forceps (Fine Science Tools: 11000-12)
- 1x scissors (Excelta: 290)
- 1x angled scissors (Fine Science Tools: 15010-10)
- 1x spatula (Fine Science Tools: 10089-11)
- 1x syringe pump (Harvard Apparatus Model 11 Plus)
- 1x 20 mL glass scintillation vial (Grainger: 3LDT2)
- Aluminum foil
- Tape
- Liquid absorbent mats
- Isoflurane chamber (E-Z Systems: EZ-178)
- Nose cone (E-Z Systems: EZ-103A)
- Magnetic stir plate with stir bar (Corning PC 353 Stirrer)
- Orbital shaker (Scilogex: SK-D1807-E)
- Scale
- Weigh boats
- Pipette controller (Grainger: 49WF85)
- Serological pipette tips
- 1.5 mL opaque microcentrifuge tube (Argos Technologies: 06333-80)
- Microcentrifuge tube rack
- Pipettes
- 18-gauge needle
- Leica TCS SP8 microscope

#### Software

- MATLAB (https://www.mathworks.com/)
- FIJI (https://imagej.net/Fiji)
- neuTube (https://www.neutracing.com/)

#### Reagent setup

- 10% potassium ferrocyanide (PF) solution

○Dissolve the appropriate amount of PF in deionized water (DIW) (10 g of PF per 100 mL of DIW) to create a 10% PF solution. Use a magnetic stir bar and magnetic stir plate to dissolve potassium ferrocyanide.
- 20% hydrochloric acid (HCl) solution

○Dilute the appropriate amount of stock HCl solution in DIW (20 mL of stock HCl per 80 mL of DIW) to create 20% HCl solution. Perform this step under a fume hood.
- Potassium ferrocyanide/hydrochloric acid working solution

Mix 1 part 10% potassium ferrocyanide solution per 1 part 20% HCl solution

### Cardiac perfusion protocol and sample preparation

#### Cardiac perfusion

Begin by anesthetizing a mouse using an isoflurane chamber with 1.5 L/min of oxygen and 4.0% isoflurane. Once the mouse is anesthetized, remove it from the chamber and place its snout in a nose cone at 1.5 L/min oxygen and 1.5% isoflurane. Administer a solution of lectin-DyLight-649 (200 μL, 25% lectin-DyLight and 75% saline) via retroorbital injection (see Fig 3A,B) (47,48). Allow the solution to circulate for about 20 minutes before proceeding with the cardiac perfusion (49).

**Fig 3.**
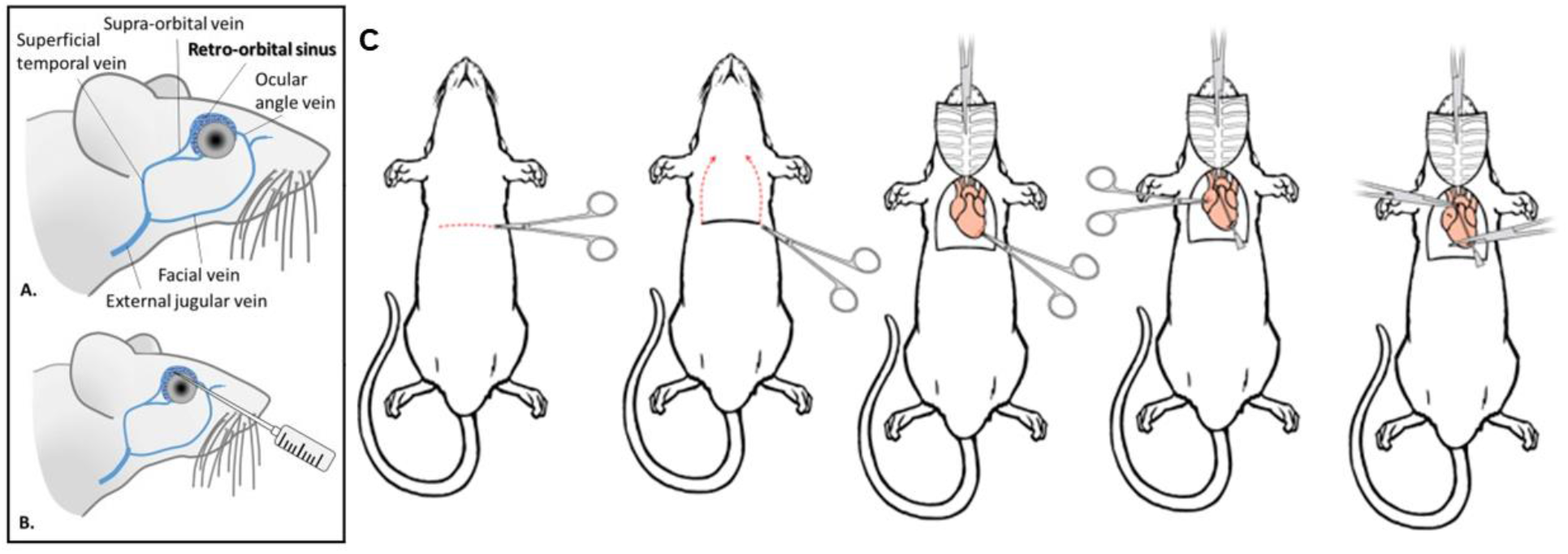
Key steps for animal procedures. A,B) Retroorbital injection diagram. Correct needle placement is shown. C) Incision points for opening the chest cavity. A lateral incision beneath the diaphragm and a vertical incision along both sides of the chest opens the chest cavity. The right atrium of the heart is cut, and a needle is inserted into the left ventricle (47,50).

It is recommended to perform the cardiac perfusion on a surgical tray or similar platform to contain the exsanguinated blood. Confirm that the mouse is at an appropriate plane of anesthesia using toe and/or tail pinches. Next, open the chest cavity by performing a horizontal incision beneath the rib cage and a vertical incision along both sides of the chest (see Fig 3C) (47). Use hemostats to assist with holding the chest open to access the heart. Then, perform a small incision on the right atrium of the heart to allow blood to exit the body. This is followed by inserting a butterfly needle into the left ventricle of the heart. Use a syringe pump to perfuse 10 mL of saline into the heart at a rate of 2 mL/min. Afterwards, perfuse 10 mL of formalin at a rate of 2 mL/min.

#### Brain extraction

Remove the head by cutting caudal of the skull with scissors (bone scissors or large scissors preferred). Cut the scalp to create two folds and expose the cranium. Using angled microscissors, gently cut upwards from the foramen magnum to approximately the location of bregma on the brain. Using a set of scissors, perform a lateral cut directly rostral to the olfactory bulbs. This cut should split the remainder of the skull down the centerline beyond bregma. Use fine tweezers or a spatula to pry open each hemisphere of the skull. Gently separate the brain from the base of the skull with a spatula. Sever any nerves connecting the brain to the skull. Place the brain in ~10 mL of 10% formalin to completely submerge the brain. After 24h, store the brain in PBS with 0.02% sodium azide at 4°C until further tissue processing.

#### (Optional) Exogenous labeling of hemosiderin with Prussian blue for cerebral microhemorrhage visualization

The following procedures are based on 1-mm thick coronal sections of a bisected brain. Volumes and times may need to be adjusted for tissues of different sizes.

Prepare a working solution of 10% w/v of potassium ferrocyanide (10 g of potassium ferrocyanide per 100 mL of DIW). Use a magnetic stir plate to mix the solution for at least 20 minutes. Next, prepare a working solution of hydrochloric acid that is 20% of stock hydrochloric acid. Mix the solutions of potassium ferrocyanide and hydrochloric acid in a 1:1 ratio. Approximately 5 mL of the mixed solution is used per sample. This solution should be prepared prior to each staining session.

Wash samples in 5 mL of DIW with shaking. Submerge each sample into 5 mL of the working potassium ferrocyanide/hydrochloric acid solution for 1 hour. Afterward, perform a final DIW wash for 5 minutes. Store the samples in PBS with 0.02% sodium azide at 4 C°.

#### Tissue clearing

The following procedure is modified from the established iDISCO protocols (4). Wash durations were modified for 1-mm thick coronal sections of a bisected brain.

Perform a series of methanol washes (20%, 40%, 60%, 80%, 100%, and 100%, balance DIW) each for 20 minutes with shaking. Microcentrifuge tubes of 1.5 mL (or larger for larger samples) are recommended. Fill the tubes fully to minimize exposure to oxygen. Next, incubate the samples in a solution of 66% dichloromethane and 33% methanol for one hour with shaking. The sample may be stored overnight in this situation if desired, without shaking. Next, incubate the sample in dichloromethane twice for 15 minutes with shaking. Lastly, store the samples in dibenzyl ether at 4°C until imaging.

#### Imaging with confocal microscopy

When imaging samples that have been cleared following the iDISCO protocol, it is recommended to image the sample while submerged in dibenzyl ether. There are various methods to safely house a sample with dibenzyl ether and protect the imaging objective. A straightforward method is to create an epoxy well on a cover glass to surround the sample and hold the dibenzyl ether.

To image the vasculature labeled with the lectin-DyLight-649, use an appropriate wavelength (633 nm can work) with an emission band of approximately 650-750 nm. To visualize microhemorrhages labeled with Prussian blue, locate regions of Prussian blue positivity using a white light source. Simultaneously collect a fluorescent image of the vasculature and a transmittance image to co-register the vascular fluorescence with cerebral microhemorrhages. This workflow along with representative images are depicted in Fig 4.

**Fig 4.**
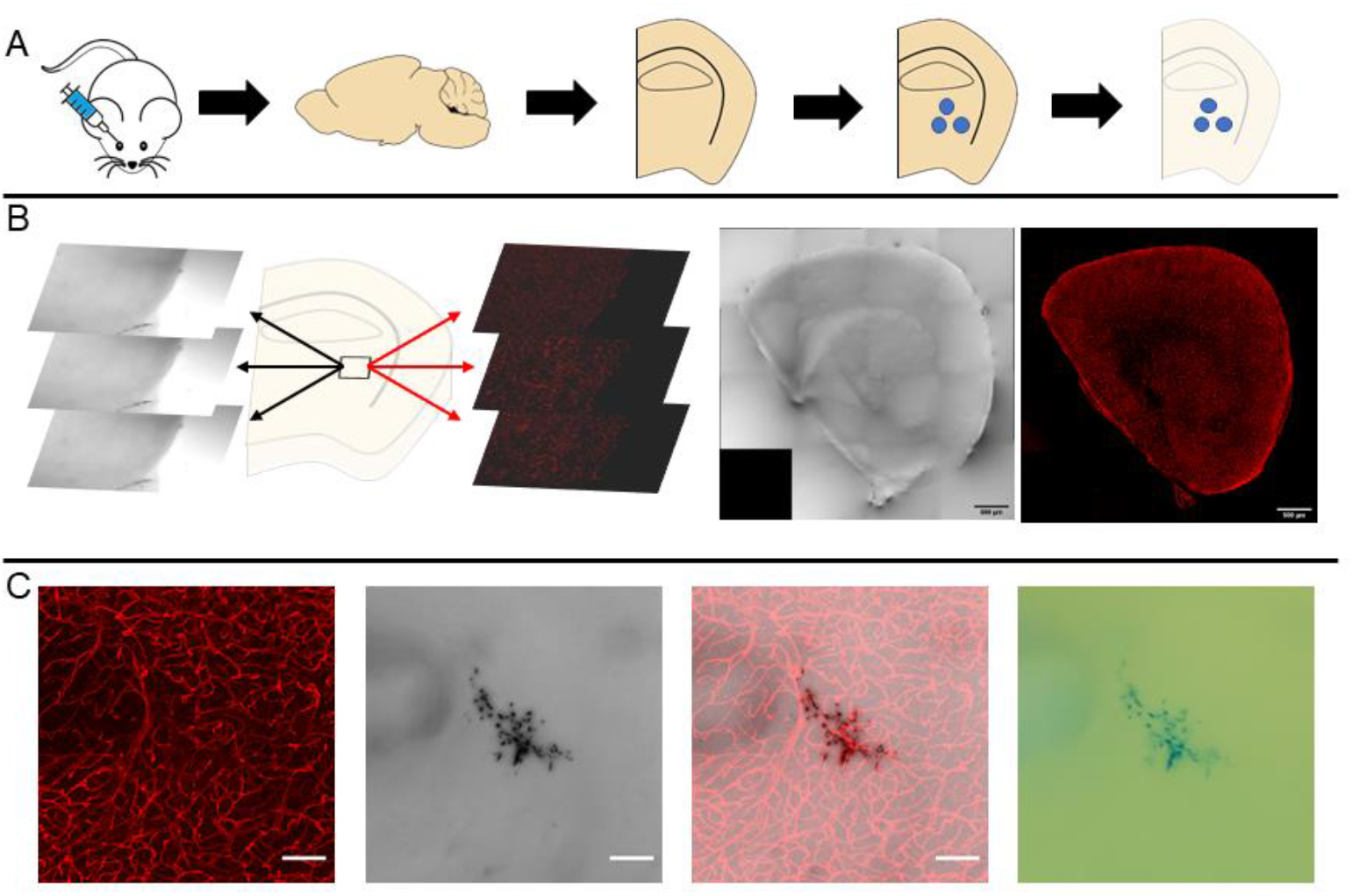
Workflow for 3D visualization of cerebral microvasculature cerebral microhemorrhage in thick tissue sections. A) Vasculature is labeled via retroorbital injection of lectin-DyLight-649. Brains are then sectioned, stained, and cleared. B) Depiction of z-stack imaging and tile stitching for a fluorescence channel and a transmission channel via confocal microscopy. C) From left to right: fluorescent image of microvascular network in a 70 μm thick tissue region, transmission image of cerebral microhemorrhage in the same tissue region, overlayed image of both imaging channels, eyepiece view of Prussian blue positive cerebral microhemorrhage. Scale bars are 100 μm.

In addition to imaging brain microvasculature, the combination of iDISCO and lectin-DyLight-649 enables visualization of the microvascular network of other organs. Light-sheet microscopy can be used as an alternative to confocal microscopy to rapidly generate 3D reconstructions of the vascular network with minimal photobleaching. Fig 5 shows representative images of different labeled and cleared organs imaged with confocal microscopy and light-sheet microscopy.

**Fig 5.**
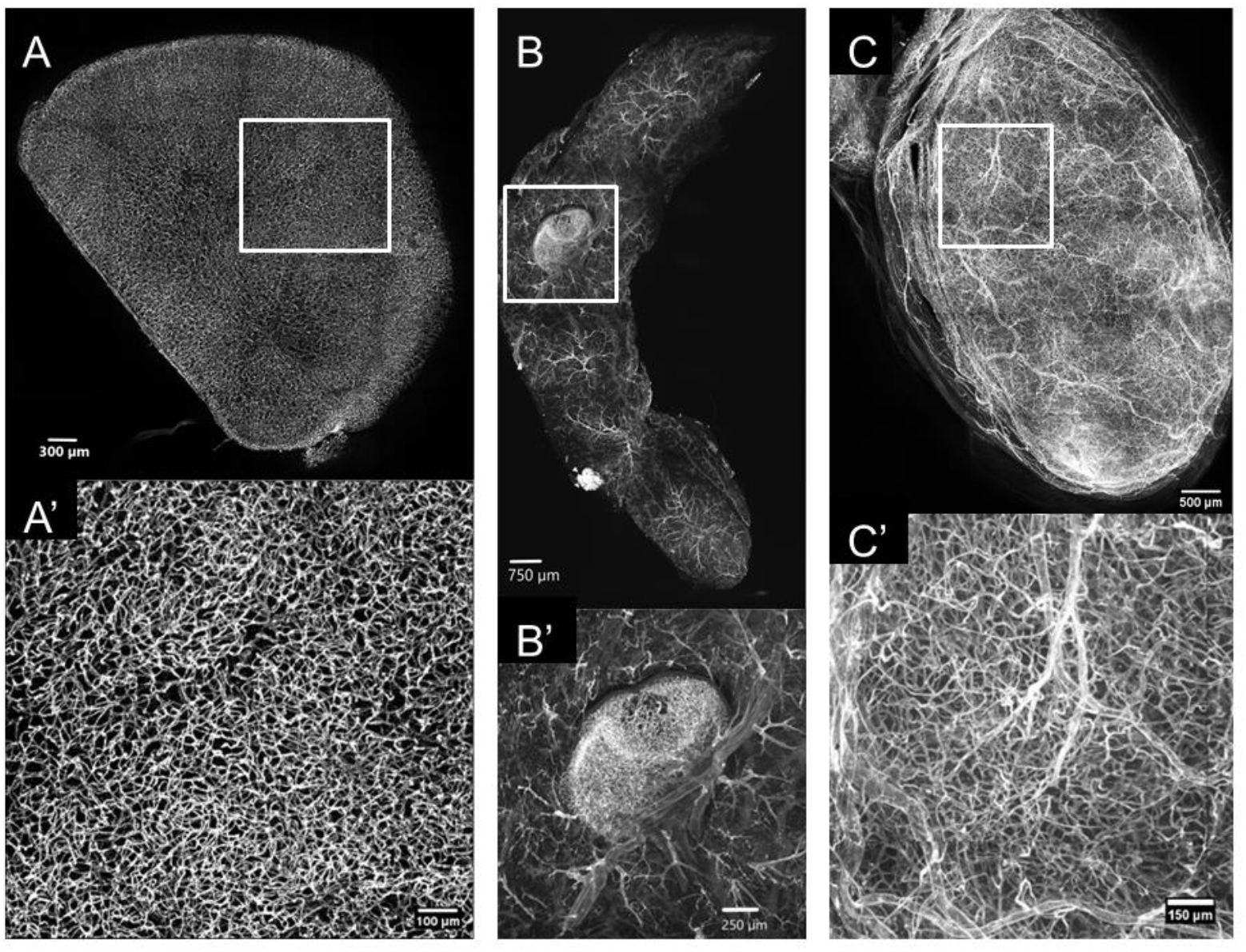
Lectin-DyLight-649 labeling and iDISCO optical clearing enable detailed visualization of mouse organ microvasculature. A) Confocal image of a 1-mm thick coronal hemisection of the brain. A’) Magnified view of A. B) Light-sheet image of a mammary gland with a lymph node. B’) Magnified view of B. C) Light-sheet image of a bladder. C’) Magnified view of C.

#### Vascular segmentation

There are various segmentation methods to isolate the fluorescent vasculature from the image background. The iterative selection thresholding method was implemented to binarize the fluorescent images (Fig 6) (51). This algorithm was selected due to its simplicity and objectivity and was performed using custom-written code in MATLAB. Vessel parameters such as vascular density and tortuosity can be calculated from the resulting segmented images. An open-source neuron tracing software known as neuTube was used to quantify vessel sizes.

**Fig 6.**
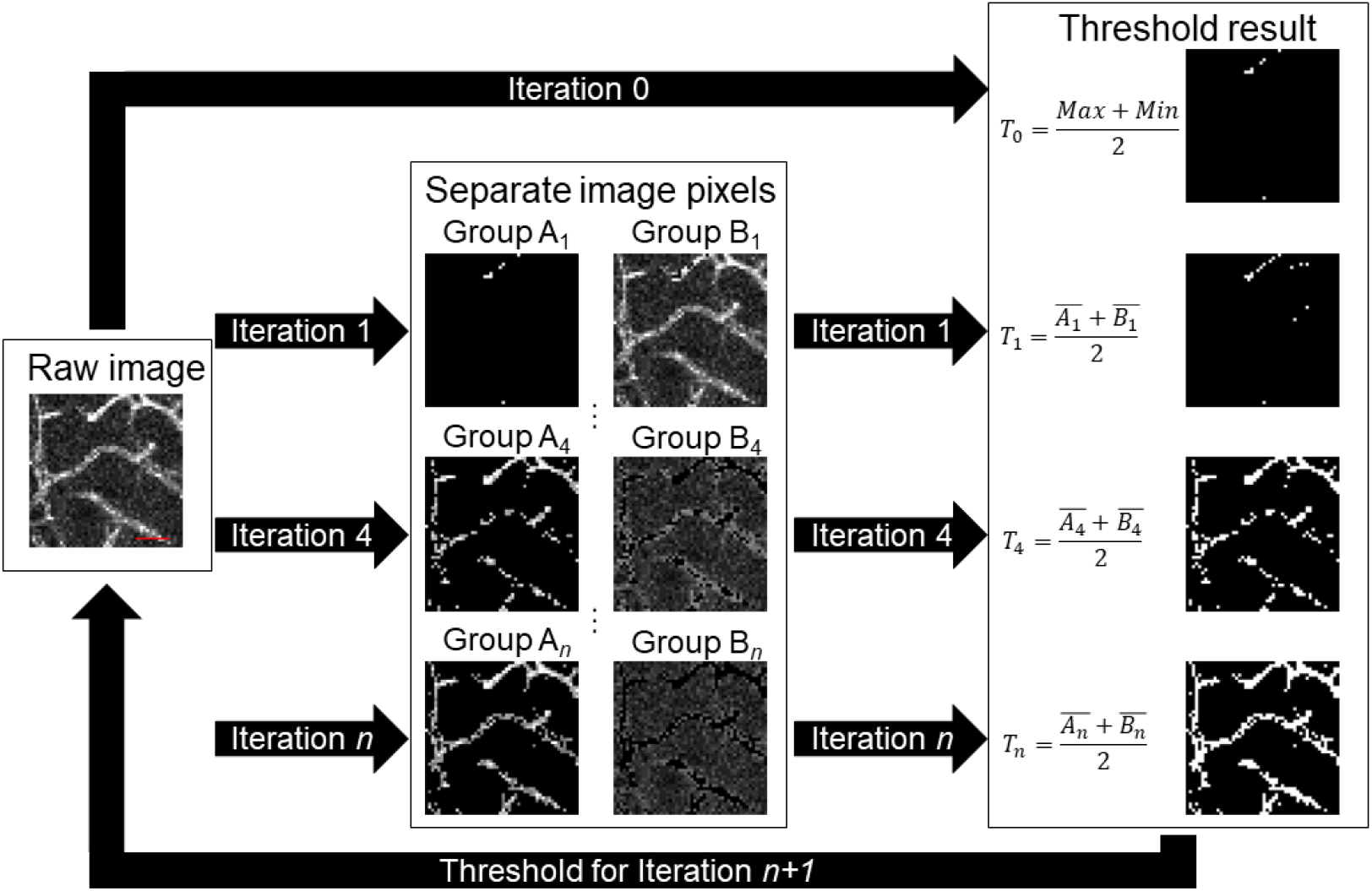
Schematic of iterative selection thresholding method for images of fluorescent vasculature. An initial threshold is selected as the average of the maximum and minimum voxel intensities present in the image. The subsequent iteration assigns voxels to group An if a voxel has an intensity greater than the threshold of the previous iteration, or to group Bn if a voxel has an intensity less than the threshold of the previous threshold. Next, the average intensity within each group is calculated, and the average of those two values determines the new threshold. This process is repeated until the change in threshold is less than one intensity value. Scale bar is 25 μm.

Using MATLAB, a 3×3×1 median filter removes noise within the image data. The iterative selection threshold method is applied by selecting an initial threshold value, *T*_0_. The initial value is the average value of the maximum intensity and the minimum intensity of all voxels of an image.

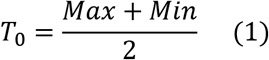

All voxels within an image are separated into two groups: group A_n_ if a voxel has an intensity equal to or greater than the threshold value of the previous iteration, or group *B_n_* if a voxel has an intensity below the threshold value of the previous iteration.

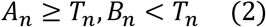

The threshold value of the subsequent iteration is calculated as the average value of the average intensity within groups *A_n_* and *B_n_*.

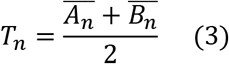

This threshold value is compared to the threshold value of the previous iteration. If the difference between the two values is less than 1, the procedure is complete, and the current threshold value is selected. If the difference between the two values is greater than or equal to 1, the procedure repeats for another iteration.

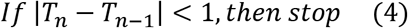

Optional morphological operations can be applied to the resulting binarized image to adjust segmentation results further. The exact parameters for these steps will vary with image acquisition parameters (particularly resolution).

For example, size filtering is performed by removing objects with a volume less than ~50 μm^3^ (the exact voxel size will vary based on image resolution) based on an 18-point connectivity scheme. Morphological closing using a spherical structural element with a radius of 1 is applied to close gaps throughout the image. Vascular density can be quantified through built-in MATLAB functions to generate a skeletonized structure, a single-pixel width line along the centerline of vessels, of the vascular image (21). From there, density can be calculated by dividing the total length of the skeleton by a given area. Vessel tortuosity can be calculated by dividing the shortest distance between two endpoints of a vessel and the total length of the vessel (52). A complete vessel is defined as the length of the vessel between two branch points.

#### neuTube tracing to quantify vessel diameters

neuTube is an open-source neuron tracing software that can be applied to tracing tubular structures, such as vasculature (46). Automated tracing can be performed on binarized vasculature images in TIF file format. neuTube will output information in an SWC format where tubular structures are simplified into individual nodes with x, y, z coordinates, a radius, and node connectivity information. Each node will be a ‘parent’ to an adjacent ‘child’ node which provides the necessary information to understand how these nodes are connected in space.

Nodal connectivity is used to identify individual vessels based on specific rules. Nodes are defined as branch nodes if they serve as the parent of two different nodes. A node is defined as an end node if it lacks a child node. Individual vessels are defined as a series of nodes bordered by two branch nodes or one branch node and an end node, including the nodes at the borders. The diameter of an individual vessel is defined as the average diameter of all nodes that make up an individual vessel.

## Anticipated results

The procedures detailed herein present a simple method for producing optically-cleared (>1-mm thick) tissue samples with fluorescently labeled vasculature. Samples can then be imaged to generate 3D images of the microvasculature. Our presented results are primarily from brain samples, but the procedures can be easily translated to other organs. The described segmentation method using custom MATLAB scripts and neuTube presents an automated method to convert the fluorescent images into binary images. The binary images can then be used to characterize microvasculature by quantifying vessel density and vessel sizes. The procedures here can be enhanced by using additional histology dyes or immunohistochemistry to label biomarkers or other structures of interest in tissue. In doing so, one can visualize and quantify the microvasculature surrounding these biomarkers to gain insight into their relationship to each other.

## Validation

### Segmentation results

Manually segmented images were used as a ground truth comparison of automated segmented images. Images from various brain sections across three different animals were used. Each section was imaged by collecting a stack of images that spans the entire thickness of the section. Automated segmentation was performed on each image stack. In addition, manual segmentation was performed on 50 randomly generated ROIs across all images. Each ROI was presented as a maximum intensity projection (MIP) image of a 50×50×5 voxel (113×113×38 μm) region. A MIP is used to present a 2D image that is more feasible for tracing. One author [DFX] was tasked with using MATLAB to outline every object within the image that the author determined to belong to a vessel. The corresponding 50×50×5 voxel region MIP from the automated segmented image was compared. Four representative examples of this are depicted in Fig 7. Across the 50 ROIs, the average sensitivity was 80±8%, and the average specificity was 90±5%. The Dice similarity coefficient between manually segmented and automated segmented images was calculated across all ROIs. This coefficient can measure similarity between two sets of Boolean data and ranges between 0 and 1, where 1 represents identical data, and 0 represents opposite data (53). The average Dice similarity coefficient across 50 ROIs was 0.71±0.07.

**Fig 7.**
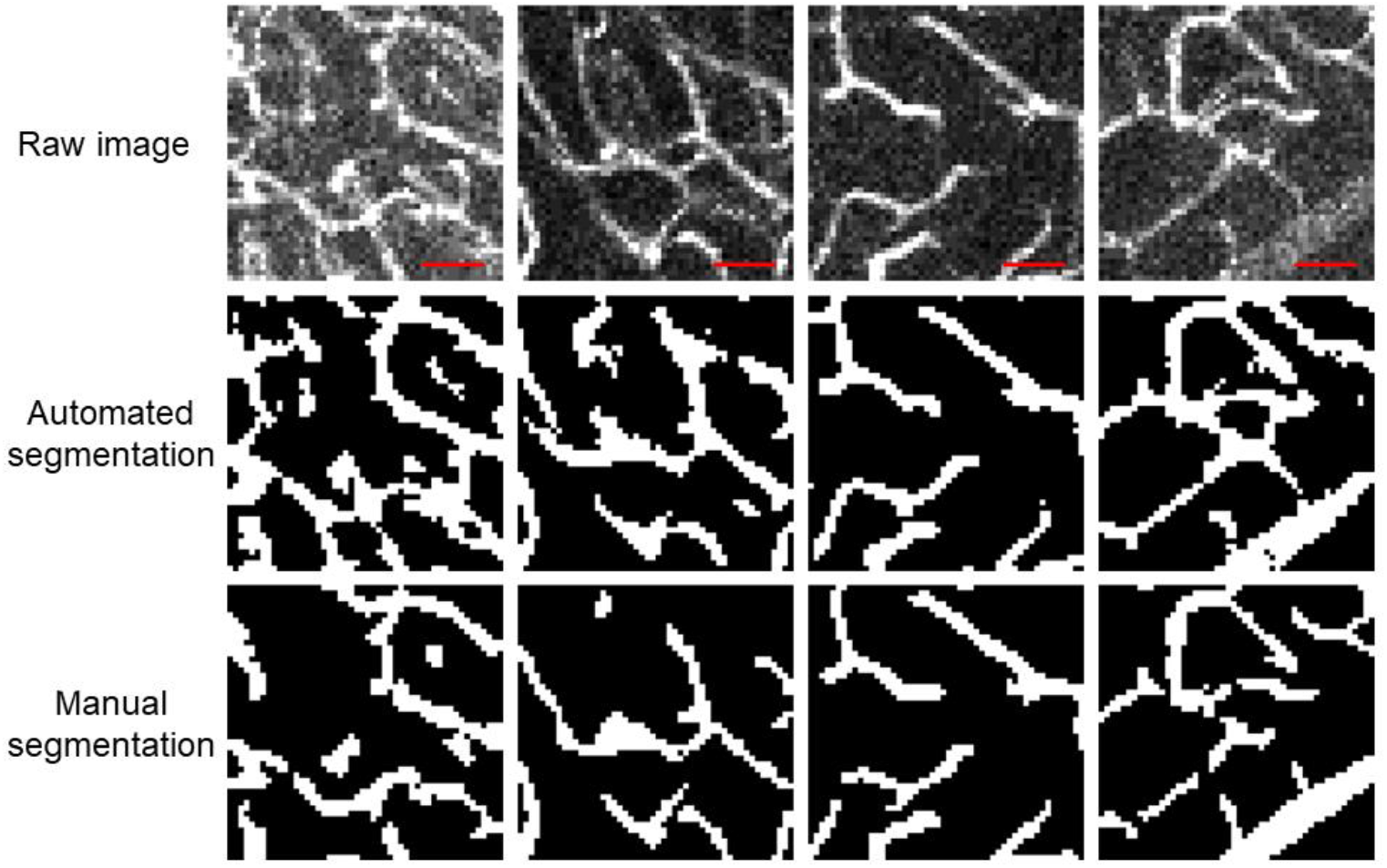
Manual and automated segmentation results. Four examples of different 50×50 pixel regions are shown. Scale bar is 25 μm.

### Diameter results with neuTube

Manually measured diameters of individual vessels were used as a ground truth comparison for neuTube approximated vessel diameters. Images from various brain sections across three different animals were used. One author (DFX) was presented with a randomly generated 50×50×5 voxel (113×113×38 μm) size region as a MIP. Within the image, the user was tasked with using MATLAB to estimate the diameter of a single vessel in the image at five different points along the vessel by drawing lines approximately perpendicular to the centerline of the vessel. In neuTube, five nodes along the corresponding vessel were selected. The average of the five manual diameter measurements via MATLAB and the five automated diameter measurements via neuTube were compared. Two representative examples of this are depicted in Fig 8. The absolute difference between diameter measurements was 1.16±0.73 μm (n=50 vessels of diameters ranging from 1.78 μm to 3.19 μm).

**Fig 8.**
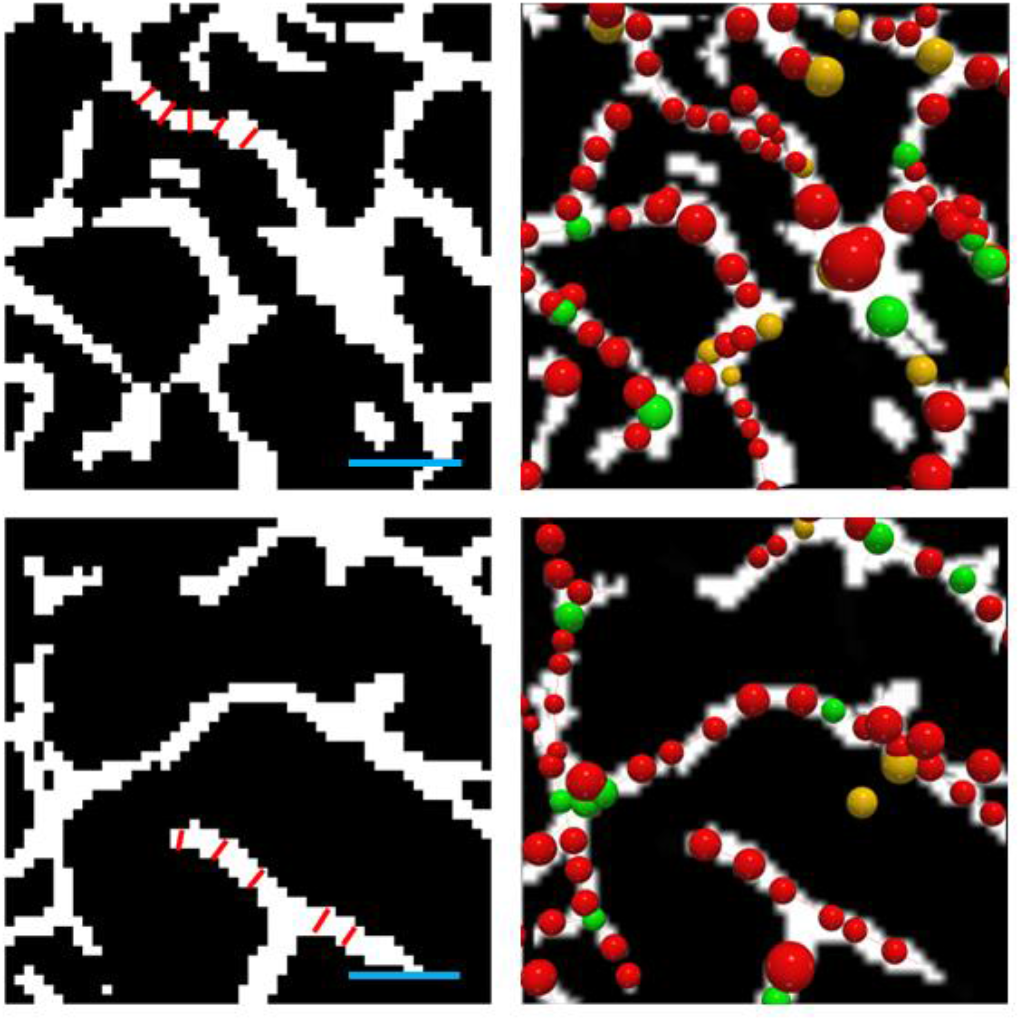
Manual and neuTube diameter results. Two different 50×50 pixel regions are shown, where one vessel is selected for manual measurement. Scale bar is 25 μm.

### Potential applications

As angiogenesis occurs in organs, endothelial cells respond to local signals to adapt vessels to the surrounding environment (54). Angiogenesis plays an essential role in tumor formation. In tumors, angiogenesis directly impacts tumor growth and metastasis. Overexpression of proangiogenic factors leads to uncontrolled vascular growth in tumors (55). As a result, anti-angiogenic drugs are frequently used as a potential treatment option for cancer. Lectin-DyLight-649 combined with iDISCO presents a robust procedure for labeling the microvasculature in all areas in the body, including tumors. Changes in vessel density, tortuosity, and diameters can be quantified to evaluate the efficacy of novel anti-angiogenic treatments.

Ischemic strokes occur when there is a significant drop in cerebral blood flow due to occlusion in a cerebral artery. A major consequence of such an event is necrosis of neurons due to deficient blood supply (56). A method to visualize the microvascular network and the surrounding neurons, astrocytes, and glial cells can provide a detailed 3D snapshot of the brain in response to an ischemic stroke and monitor potential treatments over time.

In addition to visualizing cerebral microhemorrhages within the surrounding microvascular network and providing improved capability to estimate size range of these lesions, our approach can provide enhanced imaging for other disease entities. For example, a growing body of literature suggests a contribution of dysfunctional regulation of cerebral blood flow and various types of cognitive impairment (3). The ability to visualize the cerebral microvasculature in three dimensions offers a new gateway potentially leading to the identification of novel treatment targets for neurological disorders.

## Acknowledgements

This work was supported in part by the Arnold and Mabel Beckman Foundation and the National Institutes of Health (Nos. R21AG066000, 5TL1TR001415-07, P41EB015890, R01NS020989, and R21NS111984). This study was made possible in part through access to the Optical Biology Core Facility of the Developmental Biology Center, a shared resource supported by the Cancer Center Support Grant (CA-62203) and Center for Complex Biological Systems Support Grant (GM-076516) at the University of California, Irvine

